# Coupled active systems encode emergent behavioral dynamics of the unicellular predator *Lacrymaria olor*

**DOI:** 10.1101/406595

**Authors:** Scott M. Coyle, Ellie M. Flaum, Hongquan Li, Deepak Krishnamurthy, Manu Prakash

## Abstract

Multiple active systems in a cell work together to produce sophisticated cellular behaviors such as motility and search. However, it is often unclear how this coupling specifies the complex emergent dynamics that define such behaviors. As a model system, we analyzed the hunting strategy of *Lacrymaria olor*, a unicellular predatory ciliate that uses extreme morphological changes to extend, contract and whip an apparent “cell neck” over many body lengths to capture prey. Tracking millions of unique subcellular morphologies over time revealed that these fast dynamics encode a comprehensive local search behavior apparent only at longer timescales. This hunting behavior emerges as a tug-of-war between active sub-cellular structures that use surface cilia and cortex contractility to deform the structure of the neck. The resulting search space can be described mathematically using a small number of normal shape modes that change amplitude rapidly during hunts. The distribution of these shape modes in space and time reveals a transition point between tense and compressed neck morphologies at the mean neck length, such that new shapes are readily sampled by repeatedly extending and retracting across this critical length. Molecular perturbations to the cell-signaling controller show that coupling between ciliary and contractile programs is needed to maintain this length/shape relationship; neither system alone provides the dynamic repertoire of shapes necessary for comprehensive search. Our results highlight the utility of coupling antagonistic active systems as a strategy for encoding or engineering complex behaviors in molecular machines.

**One Sentence Summary:** Analysis of millions of unique cellular morphologies of the highly dynamic single-celled predator *Lacrymaria olor* reveals that it programs a comprehensive search space and emergent hunting behavior through coupling surface based active cilia and cortex based contractile molecular systems together.

## Main Text

Cells use modular active systems to create their structure, control their mechanics and respond to their environment^1–6^. While the biochemical activities of the proteins that define these active systems and the signaling controllers that regulate activity are often known, it is less clear how cells couple these together to build up particular mechanical functions and execute specific cellular behaviors. Discovery of these basic design principles would improve our understanding of cellular function and open up new frontiers for synthetic biology and engineering at the microscale.

Within Eukaryotes, single-celled ciliated protozoa perform extraordinarily complex animal-like behaviors, such as jumping, avoidance responses, and hunting^7–11^. Because these organisms are unicellular, these behaviors must be implemented at the subcellular level utilizing multiple active molecular systems. In ciliates, two such modular systems are motile cilia patterned on the surface and contractile protein networks that operate on the cell’s interior cortex^12–17^. These systems are in turn organized in space through the geometry of the cell’s microtubule cytoskeletal scaffolding anchored to an incompressible membrane^18,19^; and in time through signaling controllers that rapidly regulate these activities^8,13,16,18,19^.

The predatory ciliate *Lacrymaria olor* is a ferocious hunter that provides an ideal model system with which to understand how higher-order cellular behaviors are built using these well-known active systems. Under a microscope, it is readily apparent that *Lacrymaria* uses sub-second extreme morphological changes to repeatedly extend and whip a slender neck-like proboscis over many body-lengths to strike and eventually engulf prey (Fig. 1A, Supplemental Movies 1 and 2). Although these dynamic morphologies appear complex, the body plan of the cell—a head, a neck, and a body—is simple enough to allow complete analysis of subcellular posture as it performs these behaviors.

**Fig. 1.**
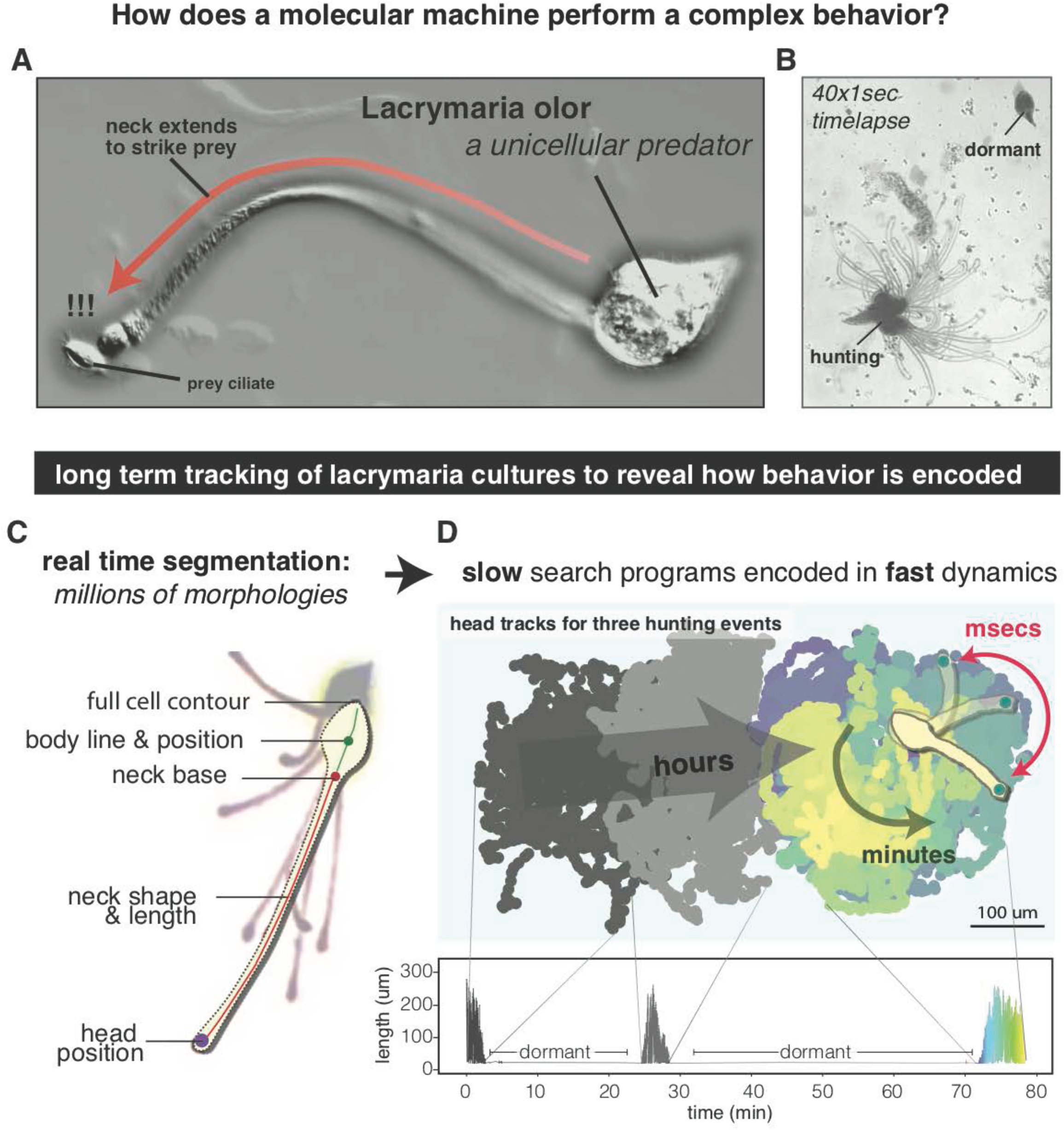
The unicellular predator *Lacrymaria olor* uses fast-timescale morphology dynamics to encode a comprehensive search behavior apparent at longer timescales. (a) Lacrymaria cell in the act of striking prey. (b) Image stack (40×1 sec) of an actively hunting cell. Although the head and neck explore local space, the body remains largely stationary. (c) Digitization of Lacrymaria morphology to track subcellular anatomy and shape. (d) Example of head location data for a single cell over the course of 80 minutes, color-coded by time. The cell sweeps out a local region of space in rapid bursts we term “hunting events” in which the neck undergoes rapid length dynamics. For this particular cell, 3 hunting events were observed and are separated by extended periods of dormancy in which the neck is fully retracted. See also Fig. S1.

We established stable cultures of *Lacrymaria* in the laboratory and used these to grow small-scale microcultures in imaging plates for long-term observation. Under these native-like conditions, *Lacrymaria* alternates between dormant states in which the neck of the organism is fully retracted; and hunting states in which the neck undergoes rapid changes in length and shape (Fig. 1B). During hunts, *Lacrymaria* attaches its tail-end loosely to the plate surface while the head, neck and body cilia are actively beating. This enables an anchored organism to use hydrodynamic forces in addition to cortex contractility to deform cell morphology. Superimposing snapshots of an actively hunting cell over time revealed that the head and neck of *Lacrymaria* sample many locations in the environment while the body remains mostly stationary (Fig. 1B). This suggested that the cell could be executing a program to comprehensively search its local environment for food.

To study this behavior quantitatively, we developed a pipeline to digitize *Lacrymaria* morphology dynamics. By incorporating techniques from computer vision^20,21^, we efficiently segmented and tracked the subcellular anatomy of individual cells in real-time movies over many hours, generating millions of unique dynamic sub-cellular morphology measurements (Fig. 1C, Supplemental Movie 3). The resulting tracks confirm that *Lacrymaria* hunting takes place in bursts of highly dynamic activity that are followed by dormant periods in which the neck is retracted (Fig. 1D). Within these active periods (henceforth “hunting events”), we find that the head uniformly samples the area surrounding its body, reaching almost all of the theoretically accessible locations with high coverage (Fig. 1D, Fig. S1). Thus, the fast-timescale morphology dynamics of *Lacrymaria* encode an emergent hunting behavior and a comprehensive search strategy only apparent at longer timescales.

We used our detailed tracking data to analyze the contributions of each subcellular anatomical structure—head, neck, and body—to building up this emergent behavior (Fig. 2A). We first examined the behavior of the fast-moving and more densely-ciliated head. Because the head is not free but attached to the end of the neck, it acts like a swimmer-on-a-string. Since the organism is also anchored to the surface, the hydrodynamic flows generated by the head motility apply direct forces to the neck and the body. To visualize the head’s velocity distribution within the reference frame of the neck, we oriented its swim velocity to the most distal segment (Fig. 2B). This distribution was asymmetric and biased towards motion oriented in line with the neck (forward/reverse) as compared to normal to it (left/right), and this bias was more pronounced when the head was moving in the backwards direction compared to the forward direction (Fig. 2B). High-speed imaging shows that the dense head cilia are highly active during extension and generate strong flows in the surroundings, but are completely inactive during retraction (Fig. 2B, Supplemental Movies 4 and 5 and 6). These results imply that when ON the head provides a strong *thrust activity* to pull on the neck in the forward direction but is turned OFF during neck retraction.

**Fig. 2.**
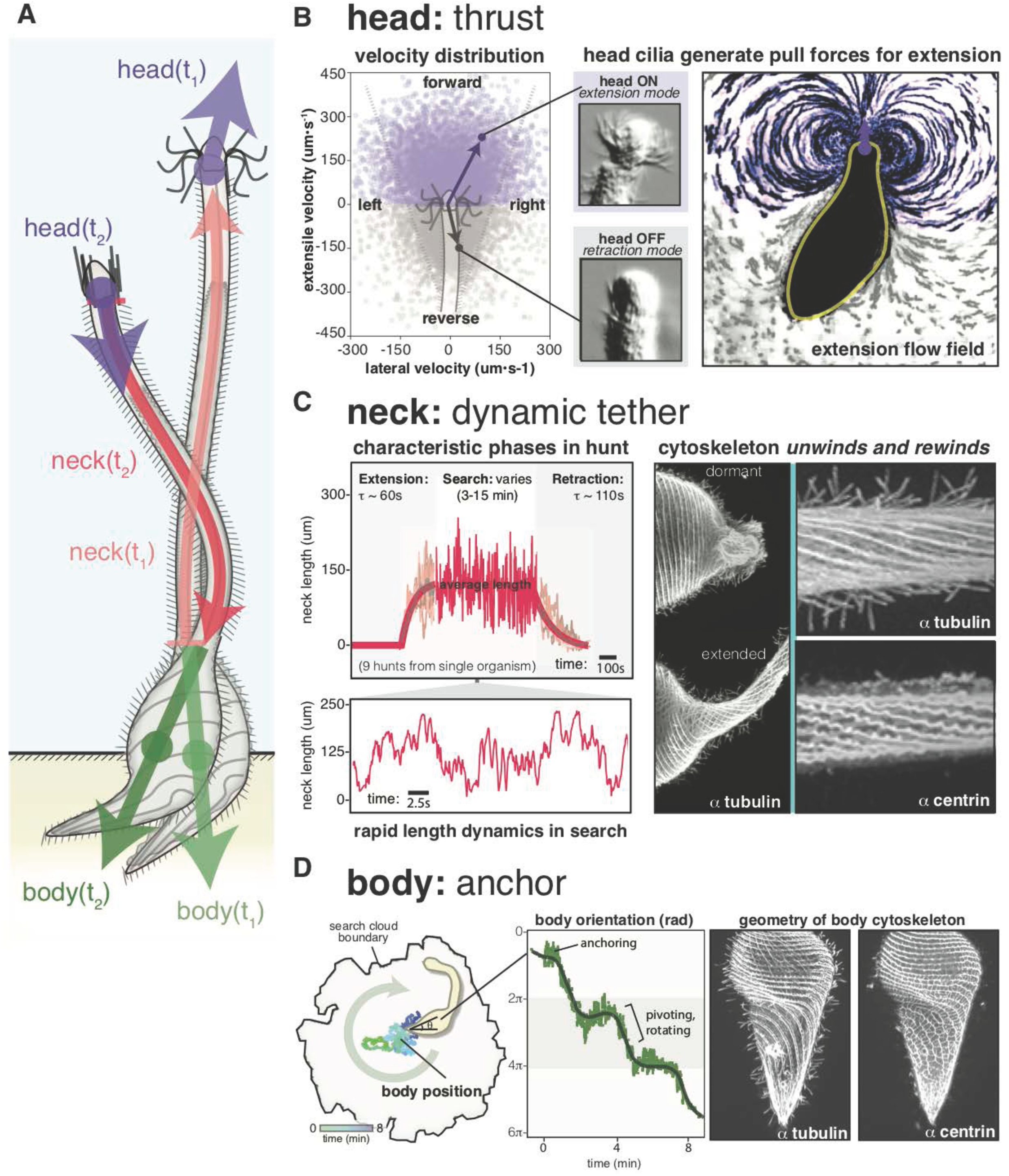
*Lacrymaria* hunting behavior emerges as tug of war between active subcellular structures. (a) Depiction of a Lacrymaria cell extending and retracting its neck. The head, neck and body segments that are tracked and analyzed are indicated. (b) Head behavior: representative velocity distribution for the head in the reference frame of the cell neck; a schematic of this reference frame is included to orient the reader. Still images taken from high-speed video show head cilia during neck extension (head ON) and neck retraction (head OFF) (see also supplemental movie 4) are shown. The stokeslet flow around the cell’s head during extension is shown by the motion of tracer particles in the cell’s surroundings as visualized by FlowTrace. (c) Neck behavior: alignment of 9 separate hunting events from the same cell with the characteristic phases of extension, search, and retraction indicated. Representative neck-length microdynamics for the search are shown below. Tubulin immunostains of a cell in the dormant versus extended states and close-up two-color immunostains for tubulin and centrin in the neck of an extended organism. (d) Body behavior: example of body centroid positions during a single hunting event, color-coded by time; body orientation for the same hunting event. Two-color tubulin and centrin immunostains of the cell body and foot.

We then inspected the behavior of the neck, which changes length dramatically throughout hunting events. By comparing these length dynamics across nine separate hunting events from the exact same cell, we found that although there was variation in the duration of hunts, there were reproducible characteristic phases common to each event (Fig. 2C, Supplemental Movie 6). Each hunt was initiated by an *extension* period (t ∼ 60s) in which the neck stretches to a mean neck length that was largely consistent across hunts; a highly dynamic searching period (3-15 min) in which the neck length rapidly fluctuates about this mean length (Fig. 2C); and a *retraction* period (t∼110s) in which the neck length decays to zero and the organism enters the dormant state. The within-organism variability in the duration of hunts that we observe was partially explained by prey capture, as most successful hunting events quickly terminated immediately after engulfment (Fig. S1). Our observations and timescales are consistent with a previous report identifying four distinct states of cellular activity in *Lacrymaria* cultures^22^.

During active search the neck may undergo hundreds of extension/retraction events of varying magnitudes and intensities. The cytoskeleton of *Lacrymaria* is reported to be arranged in a helical structure, such that these length changes do not come from a dynamic growth or disassembly process of the cytoskeleton but from winding and unwinding of the neck from this preexisting spool instead^22–24^. As such, since the membrane surface area and cell volume is conserved during this fast process, this will impose further constraints on neck deformation. We visualized this geometry by immunostaining for tubulin and centrin cytoskeletal proteins in extended and retracted cells (Fig. 2C, Fig. S2, Supplemental Movies 7 and 8). This confirmed a large helical pitch arrangement of the microtubules and cilia in the neck compared to a tightly wound arrangement with a smaller helical pitch in the body (Fig. 2C, Fig. S2). Additionally, we discovered a network of undulating centrin-containing fibers juxtaposed with the microtubules (Fig. 2C, Fig. S2). Centrin-type proteins are often core components of calcium-responsive contractile systems in ciliates^14–16,25^, and the wavy morphology of the observed filaments resembles that of the centrin-rich contractile fibers of the ciliate *Stentor*^17,25^. Taken together, these data imply that the neck is a *dynamic tether* that contains two active systems—ciliary and centrin elements—supported by a microtubule geometry that facilitates and constrains dynamic length changes during hunts.

Next, we examined the role of the body during hunts. In general, the body centroid moves very little within the boundary of the search cloud defined by the head (Fig. 2D). However, the orientation of the body does change during hunts through rotation (Fig. 2D), allowing the cell to pivot to search different regions of space. Thus, the body anchors the cell during hunting, providing crucial resistance that prevents the head from dragging the cell away during extension. Consistent with this, immunostaining shows that the spiral arrangement of the body cytoskeleton flattens out into a foot-like structure that directly contacts the surface (Fig. 2D, Fig. S2, Supplemental Movies 7 and 8). In live cells, the cilia in the helical region of the body apply a torque to the cell, while cilia within the foot region interact with the surface during anchoring (Supplemental Movie 9). Furthermore, centrin-containing fibers in this region change geometry and form a mesh-like cross-linked structure (Fig. 2D, Fig. S2); this centrin patterning has also been seen in posterior base of the contractile fiber networks of *Stentor*^25^. Thus, this geometry of ciliary, microtubular, and centrin elements likely contributes to the body’s observed function as an *anchor* during hunting events.

Our sub-cellular posture analysis demonstrates that the comprehensive local search behavior we observe during hunting events emerges as a tug-of-war between active ciliated and cytoskeletal structures that extend, retract, and deform the neck. For this strategy to be effective, the neck must be stiff enough to accommodate these applied forces, but flexible enough for it to adopt the shapes necessary to explore the local environment effectively. To characterize the flexibility and shape dynamics of the neck, we performed principle component analysis on coordinate-free length-free parameterizations of our shape data to identify the natural *eigenshape* modes of the neck (Fig. 3A)^26^. Remarkably, we found that these shapes form a series of normal harmonic modes similar to those seen for a static beam under load (Fig. 3A, Supplemental Movie 10)^27^; this is appropriate given that, irrespective of length, any neck we observe is indeed a slender object under applied forces. The first four eigenshape modes shown explain >98% of the shape variance; 90% being explained by the first two shapes alone. The fact that only a small number of shape modes is required to describe neck shape is valuable, as it permits a natural low-dimensional description of the observed dynamics in terms of the contributions of each eigenshape mode (Fig. 3A).

**Fig. 3.**
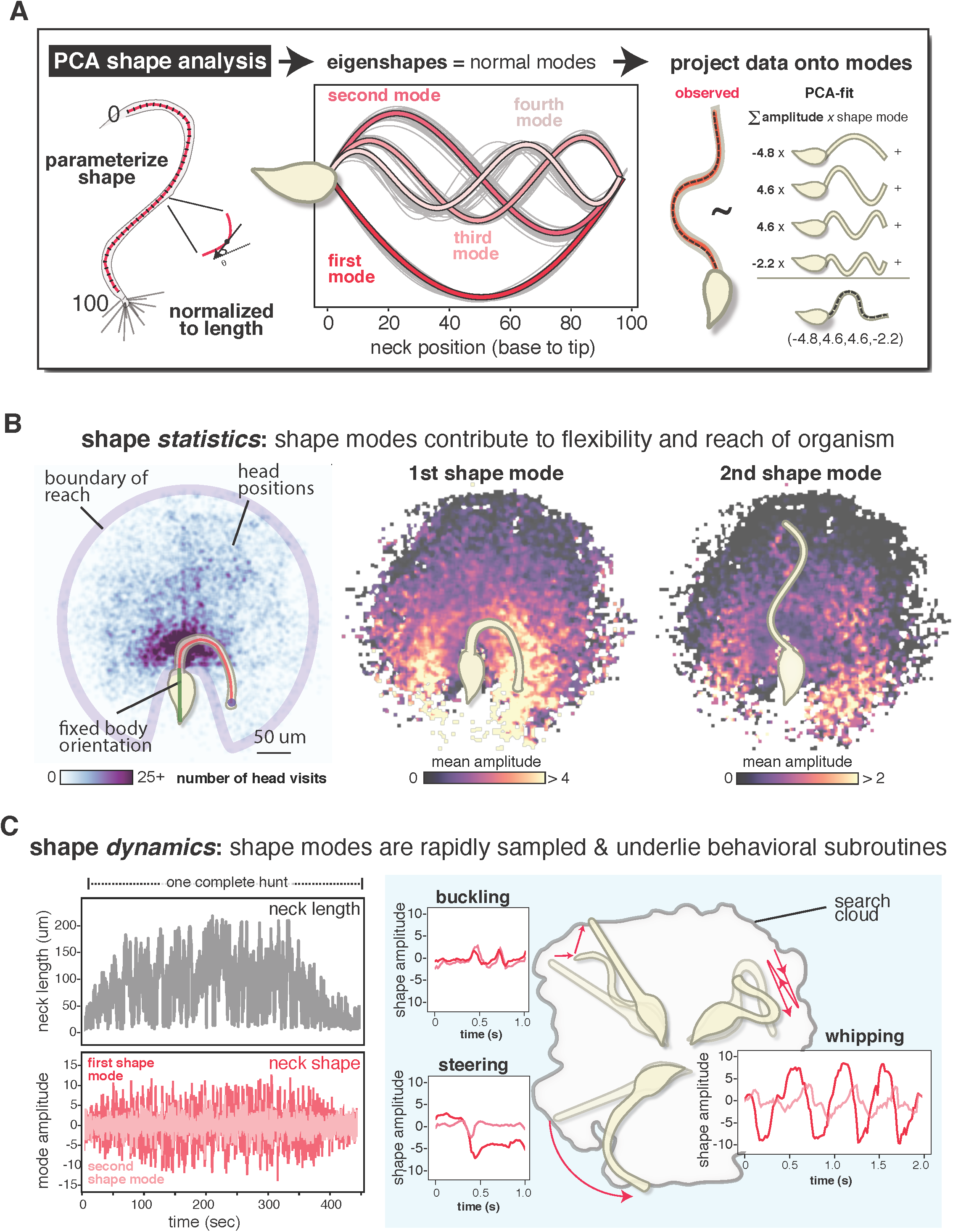
*Lacrymaria* search space can be described with a small number of normal shape modes that are rapidly sampled to facilitate reach. (a) PCA shape analysis was performed on coordinate-free length-independent parameterizations of shape. The first four eigenshapes obtained from 38 separate cells (color lines) as well as the eigenshapes from each individual cell (grey lines) are shown. Any observed neck shape can be described as a linear combination of these four eigenshapes. (b) Shape statistics: a reach plot for an organism in which all its head points are aligned to a fixed body reference frame (details in Fig. S3). This plot is recolored by the average amplitude of the first or second eigenshape mode of the neck whenever the head visits that point, showing how search space depends on accessing these shape modes. (c) Shape dynamics: necklength dynamics during a hunting event for the cell in (b) and the associated rapid dynamics of the first and second eigenshape modes. Examples of intuitive neck behaviors from experimental data–buckling, steering, whipping–that can be described in terms of patterns of shape mode dynamics are shown.

As our method of identifying these shape modes was independent of spatial coordinate, length, and time, we wondered how modes would be distributed along these axes during active search. Aligning the observed head positions from an organism to a fixed body position and orientation allows us to visualize the cell’s *reach* from its own reference frame in terms of a density plot (Fig. 3B, Fig. S3), and shows that the head can access most nearby points without moving its body. By coloring each point in this space by the mean amplitude of a given shape mode whenever the head visits that point, we find that many locations within the organism’s reach are strongly correlated to contributions from specific shape modes. For example, large amplitudes of the first mode are associated with the cell searching locations to its left or right (Fig. 3B).

The amplitudes of these shape modes undergo fast dynamics that mirror the rapid neck-length dynamics, indicating that the cell rapidly samples shape space during hunting events (Fig. 3C, Supplemental Movie 10). Unlike other dynamic cellular structures that have been previously analyzed, such as motile cilia^28,29^, these neck shape dynamics are not periodic. Nevertheless, close inspection of shape mode dynamics revealed behavioral subroutines that correspond to intuitive motions that the cell uses to sample space during hunts (Fig. 3C). These include low mode *steering* that facilitates lateral reach; low mode *buckling* that helps extended necks reorient; and high mode *whipping events* in which brief oscillations in shape modes cause the neck to wave back and forth as it rapidly reduces its length.

Since the shape dynamics are not periodic, how does *Lacrymaria* program shape-space? The physical constraints of *Lacrymaria’s* finite helical cytoskeleton are such that forces that act to extend or retract the neck may create tension or compression that leads to shape change. We examined the empirical relationship between neck length and shape mode amplitudes in our data (Fig. 4A, Fig. S4) and identified two distinct regimes: low modes at long neck lengths and high modes at short neck lengths. Intriguingly, the transition between these regimes appeared to occur at the *mean neck length* of the organism (Fig. 4A, Fig. S4). This mean length is set by the cell during the initial slow “extension” phase prior to active search (Fig. 2C), thus providing a natural transition point between compressed and tense morphologies of the neck during hunting. Indeed, when we superimposed length and shape dynamics we found that dipping *below* this mean length causes a transient spike in shape mode amplitudes as the neck buckles and compresses, while going *above* the mean length leads to a reduction in shape mode amplitudes as the neck straightens back out (Fig. 4A, Fig. S4). As such, this length-shape relationship can be exploited by the cell to rapidly and comprehensively sample shape space through repeated extension and retraction relative to the critical mean neck length that it sets during the initial phase of its hunt.

**Fig. 4.**
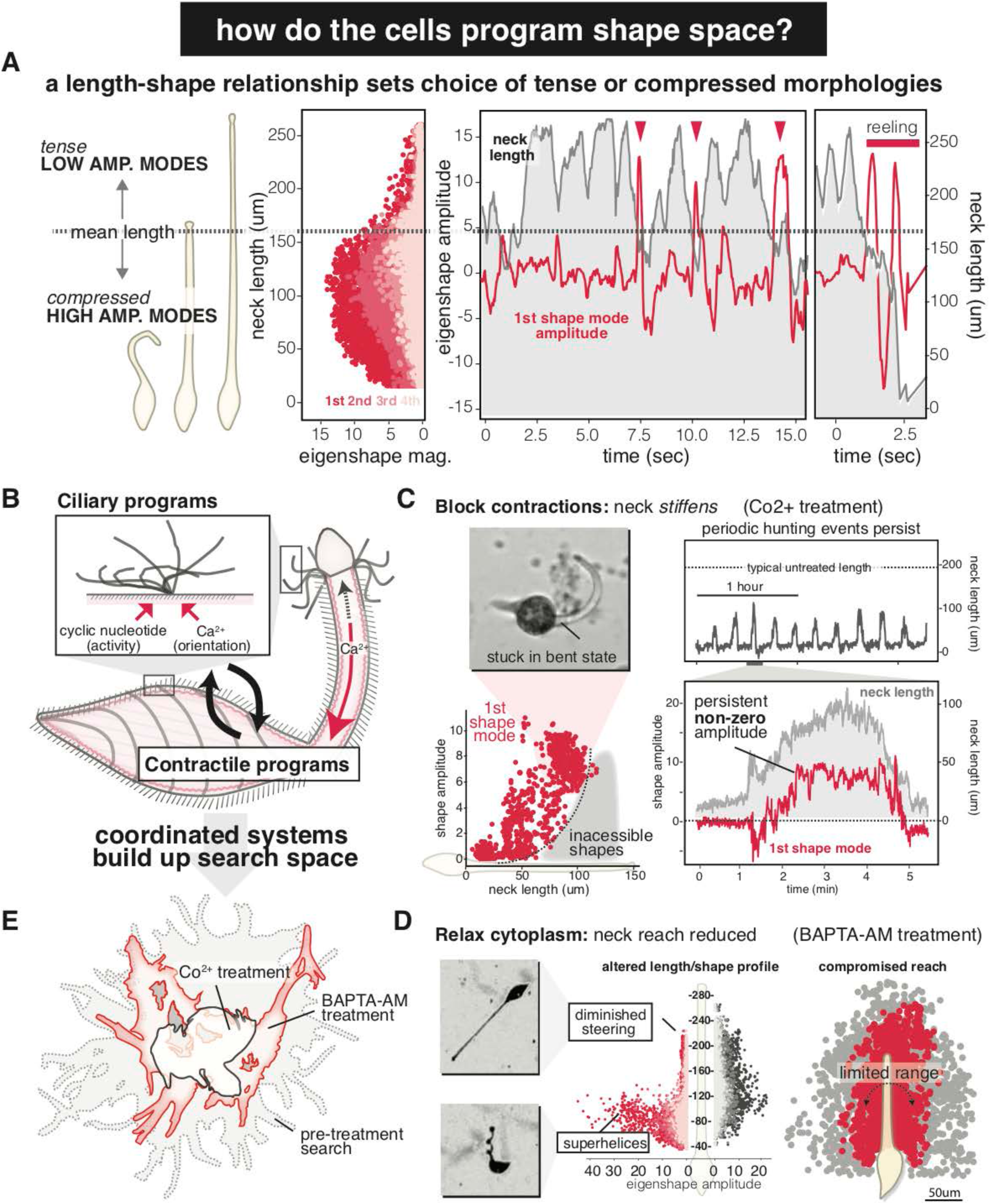
Coupling of ciliary and contractile systems is required for comprehensive sampling of shape space and to program emergent cellular hunting behavior. (a) Cartoon depiction of low (tensive) and high (compressive) shape mode regimes relative to the mean neck-length for the organism during that hunting event (dashed line) and a representative length/shape distribution for an individual cell in which each point represents an observed neck with the indicated length and eigenshape magnitude (see also Fig. S4). The associated dynamics of the neck length (grey) and the amplitude of the first eigenshape mode (red) are shown superimposed. Arrows indicate a transition to high amplitude shapes when the neck length dips below the mean length; a whipping event that reels in the neck is indicated by a red bar. (b) Schema depicting the overall architecture of the signaling controllers that regulate ciliary and contractile programs based on the data in Fig. S4 and by analogy to other ciliates. (c) Effects of blocking contractility (cobalt treatment) on hunting behavior. Periodic hunting events continue for hours in the long time-scale dynamics of a single treated cell. Neck length and shape (first mode) dynamics during these events are greatly attenuated, and the cell accumulates a persistent non-zero shape. The length/shape relationship for Co2+-treated cells is compromised and does not sample normally accessible regions. An image of a non-contractile, treated cell stuck in this bent state is shown. (d) Effect of relaxing the cell interior on hunting behavior. The length/shape relationship for the same cell pre and post BAPTA-AM treatment shows diminished steering modes at long lengths and super-helical buckling modes at short lengths. This results in reduced flexibility and diminished reach compared to untreated cells. (e) Search clouds of a cobalt-treated, BAPTA-AM treated, and untreated cells reveal how coupling between ciliary and contractile systems is required to produce *Lacrymaria’s* emergent comprehensive hunting behavior.

While this reduced-order description of the cell neck resembles that of an ordinary beam under load, the neck is a much more complex dynamic object in which length, shape, and material properties (stiffness) actively change owing to the tug-of-war of activities outlined earlier. This predicts that disrupting how these activities are coupled will strongly affect how *Lacrymaria* samples shape space and impact its overall search behavior. In other ciliates, calcium ions regulate both ciliary activity and contractility^13,16,30,31^, providing a regulatory link between active systems that act on the cell’s surroundings to those that act on the cell’s interior. Perfusion of Lacrymaria cultures with media containing elevated calcium led to increased ciliary activity and hyperextended supercoiled neck geometries (Fig. S5, Supplemental Movie 11), while perfusion with the calcium chelator EGTA caused immediate loss of cell tension and abolished ciliary reversals (Fig. S5, Supplemental Movie 12). As in other ciliates, we find that ciliary reversals and contractions appeared to require an excitable plasma membrane, as disruption of membrane polarization with potassium chloride caused transient backwards swimming followed by abnormal ciliary behavior and loss of contractility (Supplemental Movie 13)^32–34^. These and other data (Fig. S5, Supplemental Movie 17) suggest that the fundamental controllers that regulate ciliary activity and contractility in *Lacrymaria* are calcium based and likely resemble that of other well-understood ciliates (Fig. 4B).

Using this knowledge, we screened for well-known calcium ion channel inhibitors that could specifically arrest *Lacrymaria* contractility without impacting its ability to regulate ciliary reversal and activity (Fig. S5, Table S1) and found that low concentrations of cobalt chloride could produce this phenotype (Supplemental Movie 14)^35^. Cobalt-treated cells continued to perform periodic hunting events for hours (Fig. 4C, Supplemental Movie 14), but these events lacked fast length and shape dynamics; instead both length and curvature slowly rose and fell over the duration of the hunting event (Fig. 4C). Treated cells spend the majority of their hunts in a short, stiff, highly bent state and repeatedly try to push but fail to retract the neck with ciliary reversals alone (Fig. 4C, Supplemental movie 13). Consequently, there is a dramatic change to the cell’s length-shape relationship (Fig. 4C), forbidding access to a range of low mode length/shape combinations that are normally easily accessed in untreated cells. Contractility, then, functions not only to decrease length but also to reduce the material resistance of the neck, making it easier for it to be extended and stretched.

The fact that *Lacrymaria* requires active contractility to reduce the material resistance of the neck at first seems counterintuitive. However, we hypothesized imparting the neck with such a dynamic, active stiffness could play an important role in allowing cells to effectively sample shape space. To test this model, we artificially relaxed the cell interior by treating cells with the intracellular-specific calcium chelator BAPTA-AM, disrupting calcium-dependent cytoskeletal networks while leaving extracellular-calcium regulated ciliary reversal largely unaffected (Supplemental Movie 16)^36,37^. BAPTA-AM treated cells retained ciliary reversals and fast dynamics (Fig. S6) but displayed an altered length/shape relationship compared to untreated cells: extending cells had little to no left/right steering (modes diminished), while retracting cells formed tight superhelices rather than wide buckles (modes increased) (Fig. 4D, Fig. S6). The net effect of these changes is that the BAPTA-AM relaxed cells had a reduced *reach*, sampling fewer points in their local environment than untreated cells (Fig. 4D).

Our perturbation experiments hint that unlike a passive beam, the neck appears to demonstrate highly non-linear and anisotropic stiffness properties. The neck of *Lacrymaria* is an intriguing geometrical object (helical cytoskeleton) on which coupled ciliary and contractile active systems act antagonistically. The fact that the neck extends and retracts rapidly requires cytoplasm in the neck to flow back and forth (Supplemental movie 15) which can possibly generate a gradient of viscosity across the neck, stiffening and softening during a rheological cycle. Moreover, helical geometries have been known to support twist-bend instabilities that could play a further role in contributing towards the dynamic effective stiffness^38,39^. Altogether, the resulting shape space of this active filament enables Lacrymaria to comprehensively search its surroundings and hunt effectively (Fig. 4E).

By analyzing the behavior of a complex cellular machine in action, we have clarified how sophisticated emergent programs can be encoded by antagonism between coupled active molecular systems. Our approach can be easily applied to other cellular systems to understand their actions as the output of a molecular machine. The fact that coupling of independent active systems by a common signaling currency produces emergent outputs more complex than either of the underlying systems alone suggests that other cellular behaviors might be understood in terms of this paradigm. More generally, our results suggest that the organization and control of active systems can be as powerful a tool as the activities of those systems themselves for programming biological systems and engineering molecular machines at the microscale.

## Acknowledgements

S.M.C. is a Helen Hay Whitney Fellow supported by the Helen Hay Whitey Foundation E.M.F. is supported by a Stanford Biophysics NIH training grant. H.L. and D.K. are supported by Stanford Bio-X Graduate Fellowships. This work was also supported by NSF CCC grant (DBI- 1548297) (M.P.), US Army Research Office grant (W911NF-15-1- 0358) (M.P.), CZI BioHub Investigator Program (M.P.) and the Howard Hughes Medical Institute (M.P.). We especially thank R. Yanase for providing starter cultures and protocols and R. Howie for general culturing advice. We thank members of the Prakash Lab, Weeks, A.M., Pack, L.R. and Benson B., for helpful discussions and comments.

## Supplementary Materials

To facilitate viewing supplemental movies, in addition to submitting them with this manuscript we have made them available on the CiliateWorld Youtube channel (https://www.youtube.com/channel/UCRc3j56yC9zf9wEutpCLE7g) which also contains additional videos and more than one hundred hours of *Lacrymaria* 4K resolution behavioral videos.

### Materials and Methods

#### Culture methods

The field isolates of *Lacrymaria olor* that provided us with our initial behavioral observations were originally obtained from the Palo Alto Duck Pond (37.4571° N, 122.1088° W). The large-scale cultures needed for quantitative behavioral studies were grown using a *Lacrymaria* starter culture and *Cyclidium* prey culture both provided and characterized by Ryuji Yanase of Hyogo University, Japan. Cultures were grown using methods developed by Yanase^1^, with modification to facilitate long-term behavioral imaging of small-scale cultures that we describe here.

*Cyclidium* was grow using 0.01% Knop’s solution as the liquid medium (Flinn Scientific). Low maintenance “time-release” cultures were grown in darkness in 250 mL 0.01% Knop’s on top of a 25 mL nutritional agar pad (2% Agar, x mg/ml Yeast Extract, x mg/ml Skim Milk). These cultures support a modest density of *Cyclidium* for 2-3 weeks. To improve *Cyclidium* yields for feeding to *Lacrymaria*, the contents of a 250 mL flask were transferred to 8×160mm petri dishes and expanded in the dark for 3 days. These high-density cultures were pooled and 2.5 mL of this culture was used to start new 250 mL cultures at a 1:100 dilution.

To feed *Cyclidium* preys to *Lacrymaria*, 250 mL of expanded *Cyclidium* culture was filtered first through a 70 um and then through a 40 um filter (Pluriselect). The filtered cells were harvested by centrifugation at 500xRCF, and washed twice with one culture volume of fresh 0.01% Knop’s. The resulting pellet was resuspended in 100 mL 0.01% Knop’s and provided enough food to sustain 4×250mL *Lacrymaria* cultures for 3-4 days.

Lacrymaria cultures were grown in T225 TC-treated tissue culture flasks (Corning). We found that TC-treatment improved attachment of Lacrymaria in large-scale cultures and facilitated expansion and growth. Lacrymaria were grown in a standing volume of 200 mL 0.01% Knop’s in the dark. Nutrition was provided exclusively in the form of concentrated and cleaned prey foods to prevent excess bacterial growth and maintain homogenous culture conditions. *Lacrymaria* were fed 250 mL washed and cleaned *Cyclidium* per 1000 mL of *Lacrymaria* culture twice a week.

After feeding, 25-50 mL of Lacrymaria culture was removed and the cells were concentrated at 500xRCF and resuspended in 1/20 volume of 0.01% Knop’s. 100-200 uL aliquots of the resulting cell-suspension were dispensed into the wells of a 96-well Corning imaging plate. This centrifugation initially disrupts Lacrymaria morphology, however within 24 hours most cells have repaired any damage to their necks, attached to the culture plate and perform regular hunting events. Lacrymaria subcultures grown in these plates could persist for 1-2 weeks, depending on the initial density of prey in the culture.

As the natural ecosystem of these plates depends on small bits of agar and detritus to support the base of the food chain, these subcultures were often too dirty for certain imaging applications. When cleaner cultures were required, for example for high contrast dark-field imaging, cells were carefully pipetted out of one microwell and transferred to a clean one, leaving most of the sticky particulate matter, agar, and detritus behind on the bottom of the well. These cleaned subcultures could typically grow for 2-3 days before exhausting.

#### Immunostaining and Confocal imaging

Immunostaining was performed as for other ciliates with modification to preserve three-dimensional neck shape^1,2^. Briefly, cells were grown in 96-well plates in 100 uL volumes of Knop’s media overnight and fixed for 30 minutes directly in the well by addition of 100 uL of fixative solution (2% (w/v) PFA and 0.5% (w/v) Triton-X-100 in PHEM buffer). The top 100 uL of the fixative solution was then removed and replaced with 0.05 M glycine in PBS. The cells were left in this solution for 20 minutes then treated with blocking solution (0.2% bovine serum albumin (w/v) in PBS) for 30 minutes. The cells were then incubated in antibody solution (0.1% (w/v) BSA, 0.1%(w/v) Triton-X-100 in PBS) containing a 1:500 dilution of anti-centrin mouse monoclonal antibody (Millipore, 04-1624) for 1 hour. The cells then were washed 3 times for 10 minutes in PBS containing 0.2%(w/v) Triton-X-100 and left incubating in Alexa Fluor 568 conjugated □-mouse IgG antibody (Invitrogen) diluted 1:1000 in the antibody solution for an hour. Cells were again washed 3 times for 10 minutes and left incubating in antibody solution containing the Alexa Fluor 488 conjugated □-alpha-tubulin antibody (Invitrogen) diluted 1:500 for 1 hour. The cells were washed 3 more times and then mounted onto glass slides. Images were collected using either an LSM 780 or LSM 880 microscope using the airyscan technique^3^ where indicated.

#### High-speed imaging

High-speed imaging of head and neck cilia was performed using a 40x DIC Objective (Nikon) and a Phantom high-speed camera. Cells were sealed between a slide and coverslip using 2 pieces of double-sided tape as a spacer. Confinement was necessary to effectively resolve cilia with the high-speed camera. Although cells confined in this way had reduced mobility and reach compared to cells grown in unconfined cultures, cells still performed repeated extension and retraction of their necks. Data were acquired at a framerate of 1000 frames per second using the associated Phantom software, and the resulting files were converted into ordinary AVI files for inspection and analysis.

#### Live-cell fluorescent imaging

For fluorescent live cell-imaging we used Cell Mask Orange (CMO) to stain the plasma membrane and Fluo4-AM to mark the cytoplasm. CMO staining was performed at 0.1X the manufacturer’s suggested concentration for 20 minutes, followed by 5 washes in 0.01% Knop’s to remove residual dye. Fluo4-AM loading was performed at final concentration of 1 uM in 20% Pluronic in accordance with the manufacturer’s instruction.

Originally, we had intended to use Fluo4-AM to explore calcium signaling dynamics during hunting events. However, we observed that Fluo4 signal was distributed in a constitutively bright phase in the cytoplasm. In our initial experiments with Fluo4, we noticed that its signal in the neck would often lag behind the initial neck extension. This suggested that Fluo4 could be used to track the flow of viscous cytoplasm into the neck during extension.

For the live-cell two-color fluorescent imaging in part 1 of supplemental movie 15, the cell was imaged using a spinning disc confocal at 20x magnification and frames from both channels were collected at 5ms intervals (Nikon). For the live-cell imaging in part 2 of supplemental movie 15, the cell was loaded with Fluo-4AM and embedded in 0.5% low-melting point agarose as has been done previously for Tetrahymena imaging^4^. Fluo-4 was imaged with a small amount of brightfield light to permit simultaneous visualization of the cytoplasm and overall cell morphology.

#### Flow-trace imaging of tracer particles

Cells from a dense 100 uL subculture were transferred to a clean 24-well 18mm inset glass-bottom imaging plate (Makitek). 5 um beads fluorescent or non-fluorescent beads (Polysciences) were added to the solution. For low-mag visualization of flows (see supplemental movie 5), the motion of particles was visualized using Flash-Red fluorescent beads (Polysciences). For high-mag visualization of flows, the motion of particles was determined using 5 um non-fluorescent beads (Polysciences) and a 40x DIC objective (Nikon). Because the cell morphology changes during extension and retraction it is difficult to coherently analyze every class of flow produced by the cell. However, the head generates very strong flows, particularly when it is fully extended, and these flows can be recognized as a stokeslet flow when the resulting images are analyzed using FlowTrace^5^.

#### Darkfield timelapse imaging

Cells from a dense 100 uL subculture were transferred to a clean 24-well 18mm inset glass-bottom imaging plate (Makitek) or 96-well polycarbonate bottom imaging plate (Corning). The well was covered with a coverslip and sealed with VALAP to prevent evaporation. After 24 hours, the cells had reattached the plate surface and were ready for timelapse imaging. Timelapse darkfield imaging for morphology tracking and segmentation was captured with an inverted scope (Nikon) with a 2x objective (Nikon) and ring illumination; or a custom-built imaging setup using a Leopard IMX-274 MIPI-CS camera (Leopard Imaging), non-inverted 2x objective (Mitutoyo), ring LED-illumination, and encoded into 4K H264 video using an Nvidia Jetson GPU. The latter setup facilitates collecting extremely long behavioral video that is suitable for analysis but also readily shareable with the scientific community and general public (available at the CiliateWorld Youtube channel).

#### Image processing, segmentation, tracking, and data processing

Our analysis of cellular behavior from timelapse darkfield video was performed using custom python scripts that take advantage of the OpenCV computer vision package^6^. The source code is available at the CiliateWorld channel, and we describe here the general approach used in our tracking of subcellular posture. Because the cell morphology of *Lacrymaria* (fat body, thin neck, motile head) contains multiple visually distinct elements, we found no single thresholding or filtering method could consistently segment *Lacrymaria* cells. Instead, we created a composite segmentation based on using filters that recognize different aspects of its subcellular anatomy: a dynamic MOG background subtractor to identify the small but highly motile head structure^7^; a very stringent simple threshold to locate the extremely bright bodies of Lacrymaria cells; and a canny-edge detector that was effective at identifying the neck that connects the two. Cells were identified from this composite segmentation by first defining them in terms of a body, and then locating the associated head and neck. Tracking of single cells was performed using this composite segmentation, and different features of subcellular anatomy and their relative positions to one another were used to orient the geometry of the organism. The resulting tracks contain all information necessary to reconstruct the complete shape, geometry, position, and morphology of cells during the timelapse (see supplemental video 3).

Our analysis of *Lacrymaria* shapes using principal component analysis was performed using a method previously used to characterize the locomotory behavior of *C. elegans*^8^. Briefly, each neck shape was fit with a spline of 100 points, the tangent angle at each point □(s) was calculated, and the resulting description was normalized such that the mean □(s) is 0. The resulting set of neck shapes (>100,000 per cell) is coordinate-free and length-independent. The covariance matrix for the shapes was calculated and the associated eigenvectors and eigenvalues determined using the Numpy package in python. Fits of observed neck data to the first four eigenshapes was performed by least squares fitting.

In some perturbation experiments in which necks adopted highly unusual geometries our segmentation code was unable to satisfactorily extract the geometry. In these instances, *Lacrymaria* necks were tracked manually using a WACOM pen in ImageJ. The resulting neck shapes were exported and loaded into python for analysis by the same methods as computationally tracked morphologies.

#### Perturbation experiments

Pertubation experiments were performed by exchanging the media in a well with media containing the target drug concentration. Whenever possible, drugs and small molecules were prepared directly in 0.01% Knop’s solution. AM-ester containing molecules such as BAPTA-AM were solubilized in 20% Pluronic in DMSO and serially diluted into 0.01% Knop’s to the target concentration. More specific details about perturbation treatments are included in Supplemental Table S1.

---

**Fig. S1.**
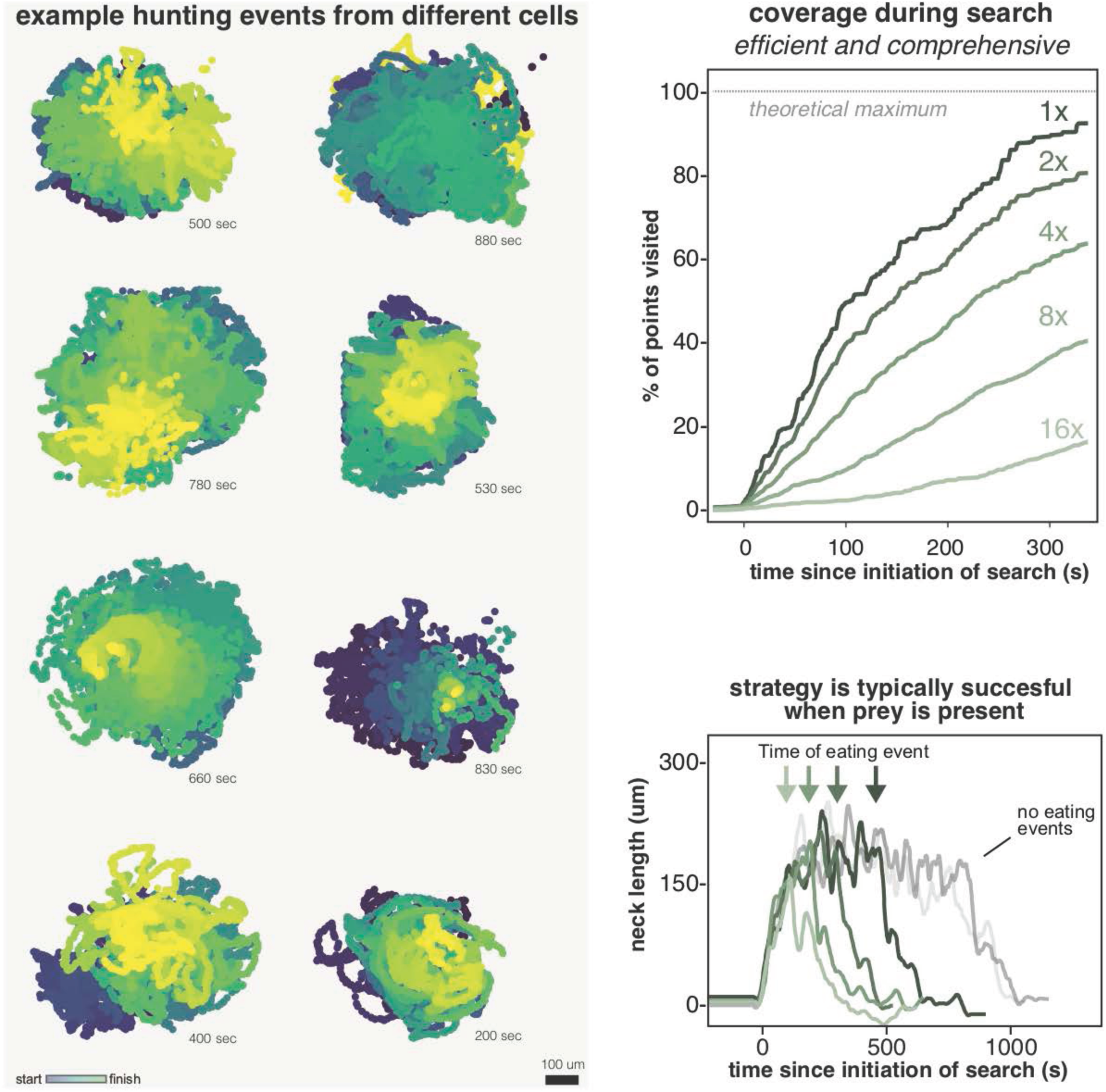
*Lacrymaria* hunting events search the local environment efficiently and comprehensively. (a) Search clouds for single hunting events from 8 separate cells. Each point in the cloud corresponds to a point the cell’s head visited and is colored by the time it occurred in the hunt (blue: start; yellow: finish). The duration of each hunting event is indicated. (b) Experimentally measured search coverage over time during a single hunting event, defined as the fraction of theoretically accessible points (based on minimal circle enclosing observed point cloud) accessed the indicated number of times by the organism. Nearly all reachable points are visited at least once, and most are visited many times over. The t_1/2_ to “fill” accessible space (∼140s) indicates that the organism efficiently samples space during search. (c) Six separate hunting events from the same cell in a culture supplied with prey organisms, aligned to the start of the hunt. Prey capture was observed in four of the six hunting events at the times indicated by the arrows, which led to retraction of the neck and a return to the dormant state. This indicates that the efficient search of space can result in effective hunting outcomes.

**Fig. S2.**
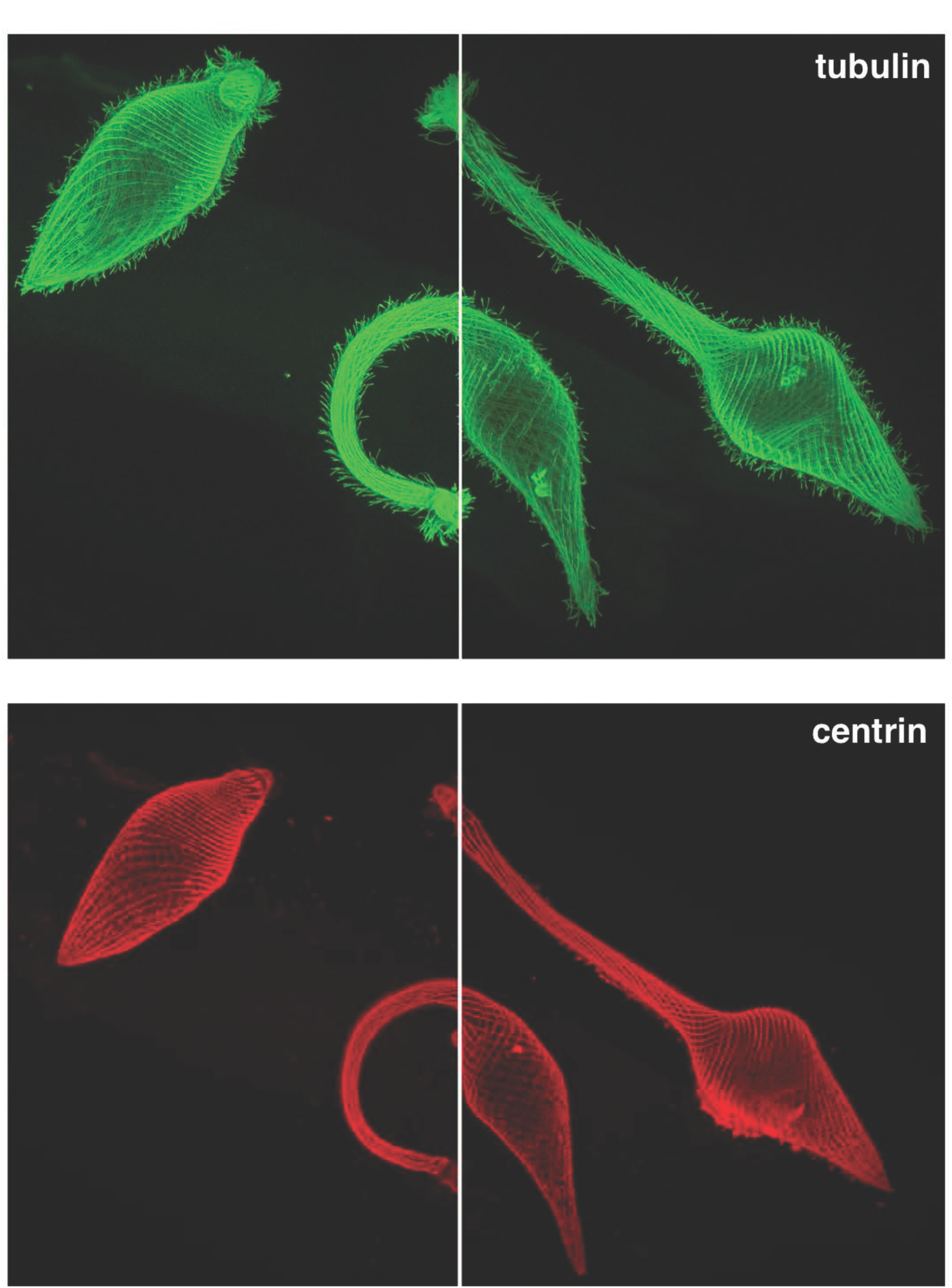
Confocal images of the cytoskeleton of *Lacrymaria* cells in different conformations. Maximum z-projections from stitching two confocal stacks of three cells in the same field of view for either (a) tubulin or (b) centrin. Additional three-dimensional representations of these cells can be found in supplemental movies 7 and 8.

**Fig. S3.**
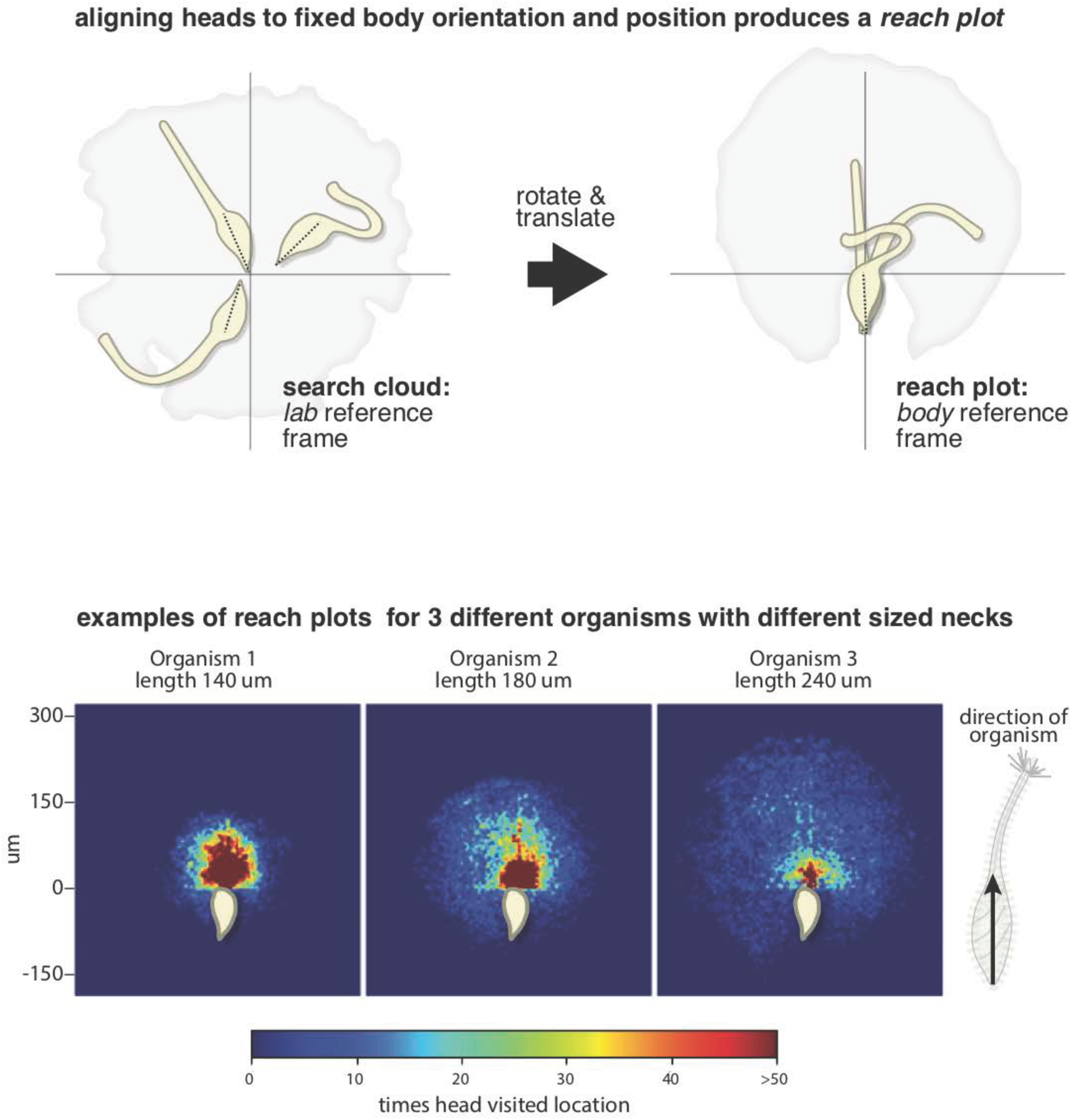
*Lacrymaria* cells have an extraordinarily flexible reach. (a) Illustration depicting how reach plots are constructed. From the lab reference frame (on the left) we observe a search cloud that the organism carves out of space. By choosing a fixed body position and orientation, we can rotate and translate each head position from our dataset to view the search from the reference frame of the cell body (on the right). Now the head positions of the head now reflect the intrinsic reach of the organism and the shapes of the neck reflect the flexibility of that structure. (b) Examples of reach density plots for three separate cells, each of which has a different size neck. The color of each point in the reach cloud indicates the frequency with which it is sampled during search. All three cells sample a wide swath of points, and can sample positions to their far left, right, and even some regions behind them. There is a clear length-dependence to these reach density plots: cells with shorter necks sample more densely much closer to them; while cells with long necks sample distant points with high frequency at the expense of less dense sampling overall.

**Fig. S4.**
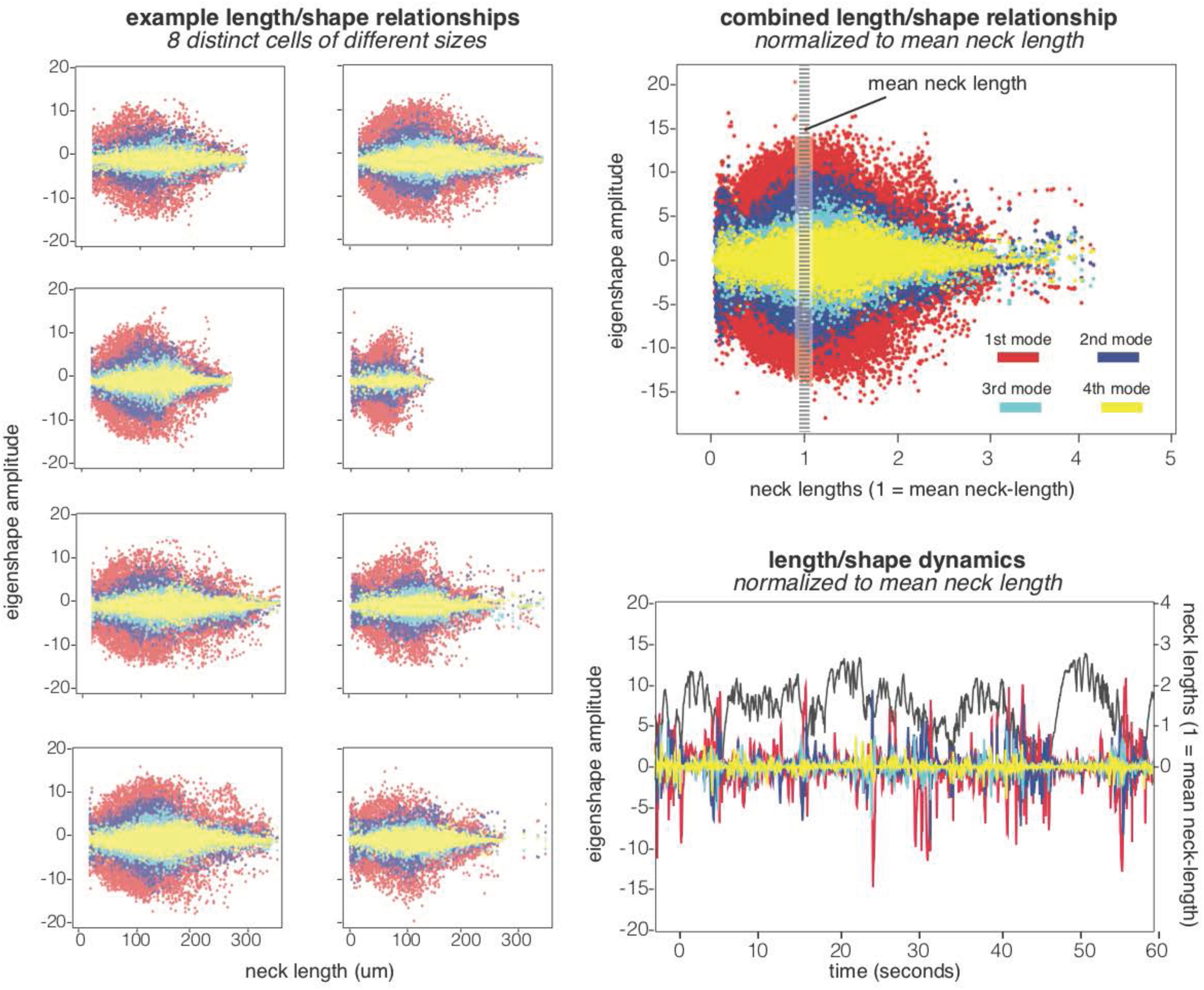
Additional examples of *Lacrymaria* length/shape relationships. (a) Length-dependence of eigenshape amplitudes (red=first mode; blue=second mode; cyan=third mode; yellow=fourth mode) for eight different cells with distinct neck sizes. (b) Aligned length/shape relationship for the eight cells in (a), in which each relationship from (a) was normalized by its mean neck length. Although individual cells vary with respect to absolute length, their length-shape relationships align very well when normalized. (c) Example of normalized length/shape dynamics (1 minute in total) for an individual cell. Note the different behavior of the shape amplitudes when the neck length is above (tense) or below (compressed) the mean neck length (length=1).

**Fig. S5.**
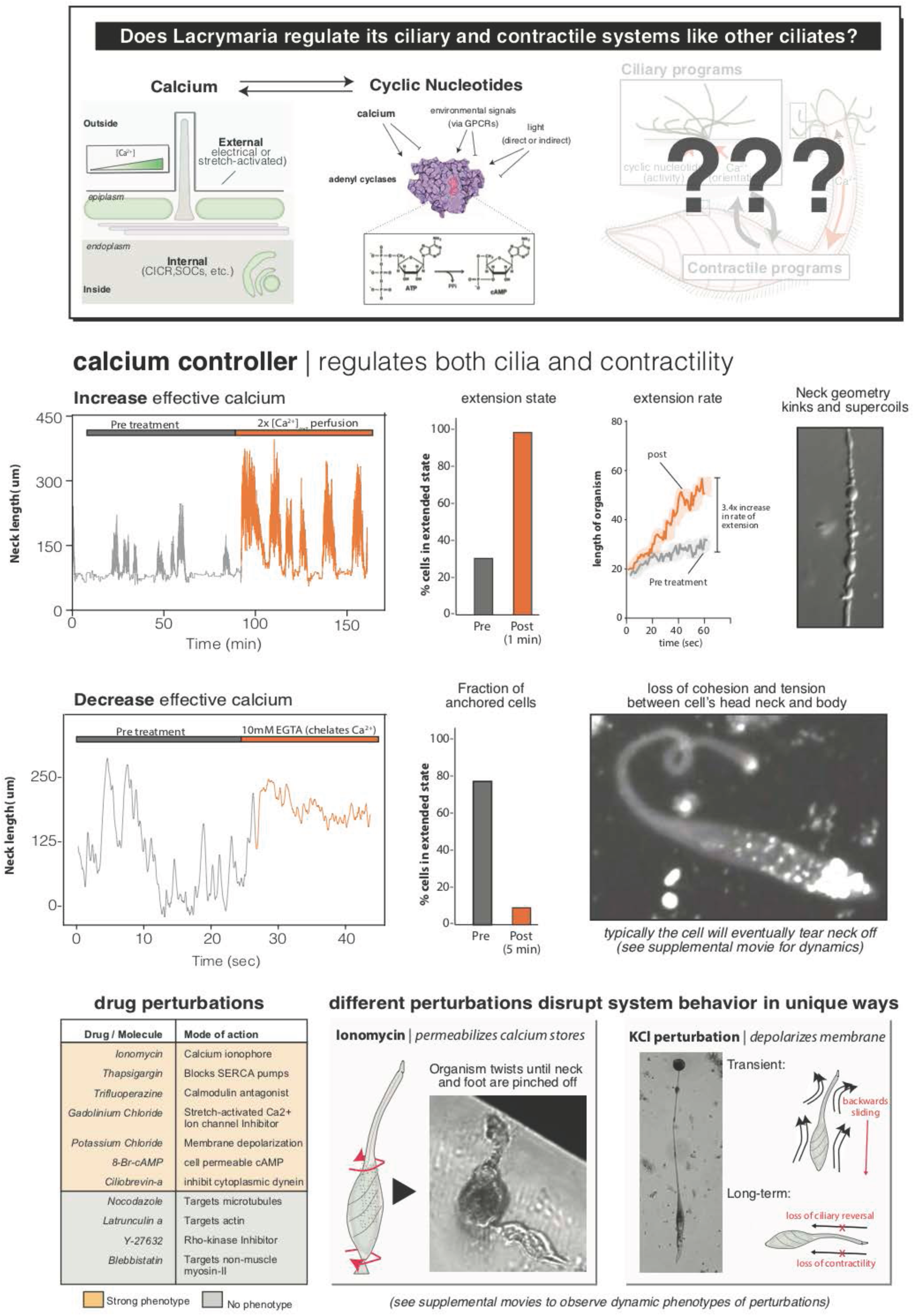
*Lacrymaria* uses a calcium controller to regulate ciliary and contractile activities. (a) Illustration depicting signaling controllers commonly used by other ciliates to regulate ciliary and contractile activities. We tested whether the main features of this system apply to Lacrymaria, acknowledging that additional genomic and molecular work will be needed to identify the specific ion channels, regulatory proteins, and factors that underlie specific details of the *Lacrymaria* controller. (b) Effect of *elevated calcium* levels on *Lacryamria* hunting behavior. A single cell is tracked for over an hour before perfusion with media containing elevated calcium at the indicated time. The cell was in a dormant state at the time of perfusion and with seconds immediately began transitioning to an active state. The cell extended to a length more than twice as long as the pre-treatment lengths, and this hyperextended length was observed in the next five post-treatment hunting events as well. (c) Inspection of the initial extension periods from the pre- and post-treatment hunting events showed that this hyperextension is manifest by a faster extension rate, rather than a longer period of extension. (d) The hyperextended form of these necks was quite prone to supercoiling and forming kinks (see Supplemental Movie 11 as well). (e) Effect of *decreased calcium* levels (EGTA-treatment) on Lacrymaria hunting behavior. Neck length dynamics before treatment show rapid length dynamics, while within seconds of treatment length dynamics are almost immediately attenuated. (f) Most cells lost the ability to stay firmly attached to the surface and began to swim freely. (g) This is coincident with a noticeable loss of apparent tension within the cells as indicated by the loose, soft appearance of the neck. Unlike the elevated calcium treatment, EGTA-treated cells cannot be tracked across multiple hunts as most cells will eventually tear the neck free from the body and cannot rebuild it unless calcium is added back to the solution. See also Supplemental Movie 12. (h) Table of drug perturbations tested related to calcium-based signaling controllers or the potential downstream targets of that regulation. Entries in the orange section of the table produced phenotypes (see supplemental movies), while those in the gray section did not—though we emphasize that we cannot confirm that those drugs work effectively in the context of *Lacrymaria* cells. (i) Examples of perturbations to calcium controller that produce distinct phenotypes owing to distinct mechanisms of action. Ionomycin, which permeabilizes calcium stores, led to immediate compaction of cells and an eventual twisting off of the junctions between the cell body and its neck and foot (see supplemental movie 17). In contrast, KCl perturbation, which affects membrane polarization, had a two-phase phenotype in which cells first transiently swam backwards before settling into a long-term state in which the cells became unable to reverse cilia and lost contractility (see Supplemental Movie 13). These observations are consistent with the notion that *Lacrymaria*, like many other ciliates, uses membrane excitability and a calcium controller to regulate its ciliary and contractile activities.

**Fig. S6.**
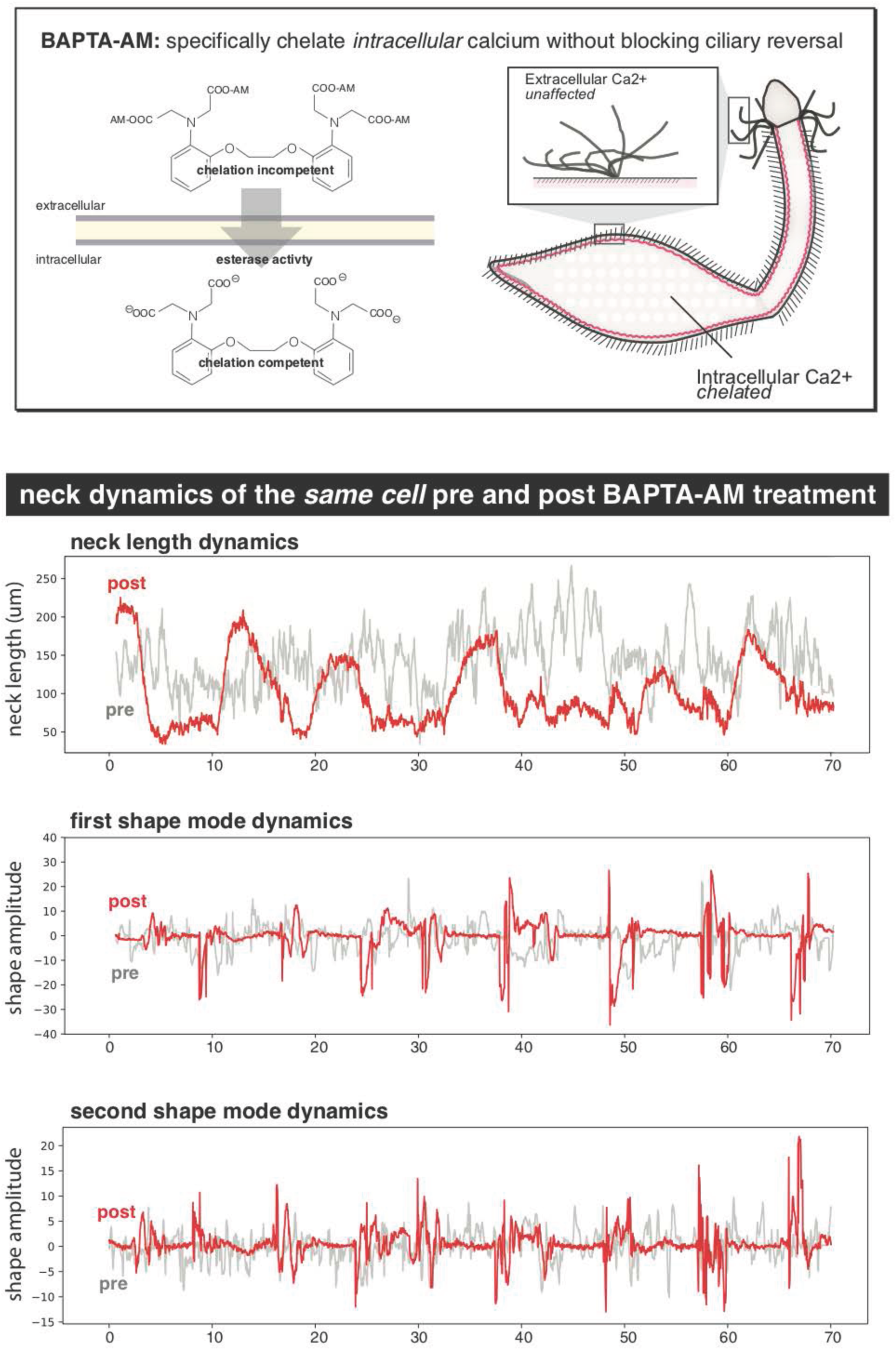
Shape and length dynamics of BAPTA-AM relaxed Lacrymaria cells. (a) Schematic showing the mechanism by which BAPTA-AM selectively chelates intracellular calcium while leaving extracellular calcium levels unaffected. The chelation activity of BAPTA-AM is restricted until the removal of the acetomethoxy (AM) esters by intracellular esterases. This allows intracellular calcium to be disrupted without affecting extracellular calcium levels, thus allowing for ciliary reversals (which depend on extracellular calcium) to be retained. (b) Neck length dynamics of the *same cell* prior to (pre) and after 20 minutes of BAPTA-AM treatment (post). The neck length spans the same range of lengths both pre- and post-treatment, however the high-frequency length dynamics are largely lost following treatment as post treatment necks monotonically extend for ∼5 sec, then monotonically retract for ∼5 sec. This indicates that the cilia are still capable applying forward and reverse forces to the organism following treatment. (c) First and (d) second shape mode dynamics for the same cell as in (b) pre- and post-treatment with BAPTA-AM. It is readily apparent that post-treated cells lack the high frequency shape dynamics of untreated organisms. Instead, treated cells undergo extended periods in which the amplitudes of all shape modes is nearly 0, followed by punctuated bursts of extreme shape amplitudes typically during the late stages of retraction.

**Fig. S7.**
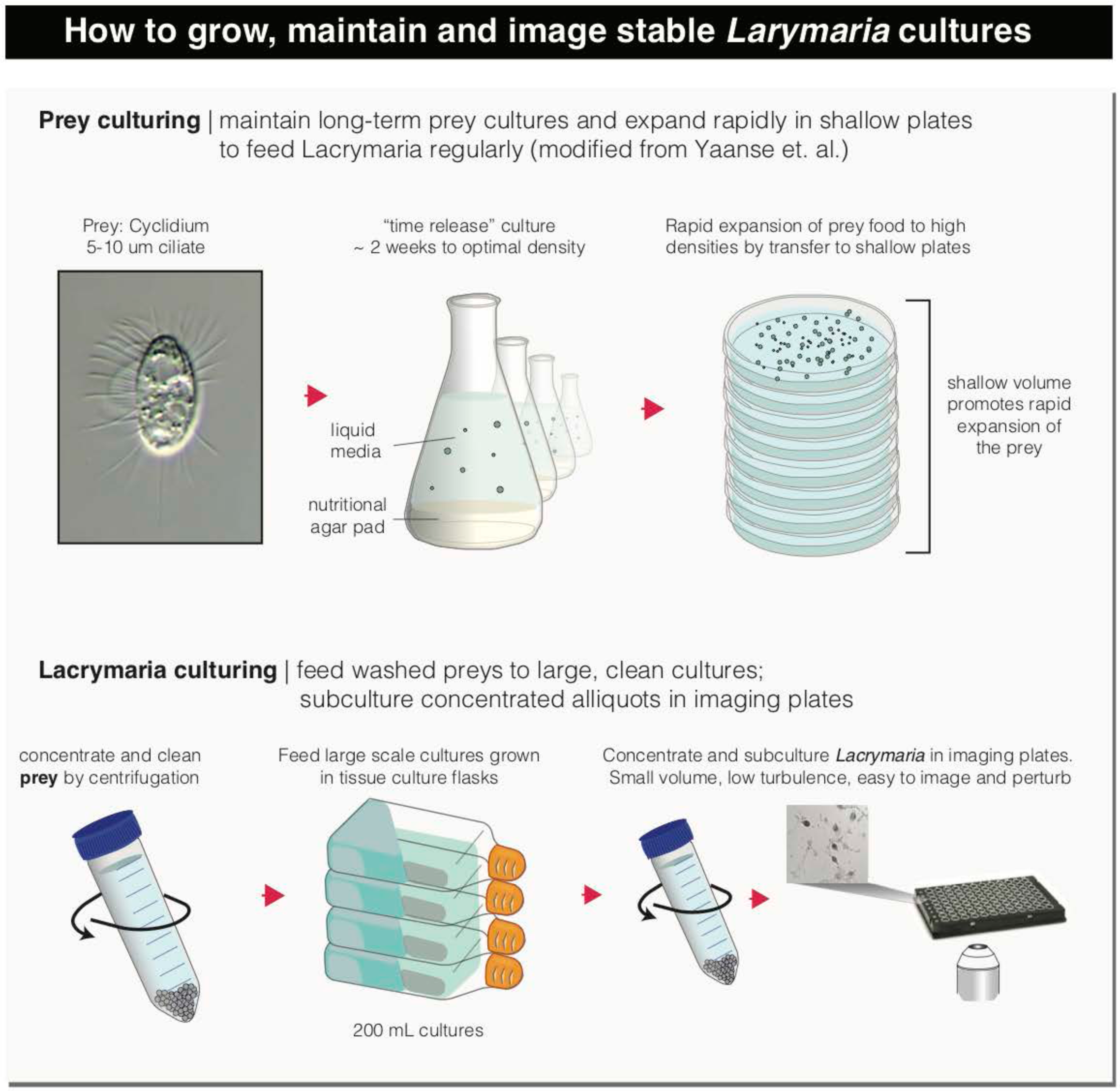
Schematic of culturing method for long-term growth, maintenance and single-cell tracking of *Lacrymaria*. Illustration depicting the process by which sufficiently dense *Lacrymaria* cultures are prepared for single-cell tracking and long-term imaging. The method of culture is adapted from Yanase et. al. with specific modification to facilitate stable growth of small-scale subcultures directly in imaging plates. Lacrymaria growth is sustained by consumption of large quantities of the small prey ciliate *Cyclidium*. These cultures are maintained at modest densities through serial propagation in liquid cultures in which the nutrition is provided through slow release from an agar pad. Transferring the liquid prey culture to 160 mm petri dishes 3-4 days prior to a Lacrymaria feeding led to a large increase in *Cyclidium* densities before the culture exhausts all available nutrition. These expanded *Cyclidium* cultures are then filtered and washed with fresh media to provide a highly concentrated, clean prey culture to feed large-scale *Lacrymaria* cultures (2x a week). Upon feeding, 25-50 mL of the large-scale culture is concentrated 25x by centrifugation and the resulting cells are dispense in 100-200 uL volume cultures in a 96 well imaging plate. Within 24 hours the majority of cells will have settled on the plate surface and perform regular hunting events. These cultures can be further cleaned for darkfield imaging by transfer to a fresh well on the plate after 24 hours to separate cells from residual agar or other debris.

**Supplemental Table S1.** List of molecules used in perturbation studies of Lacrymaria and the associated concentrations used.

| **Molecule tested** | **Concentrations used** |
| --- | --- |
| 8-Br-cAMP | 100 uM |
| Blebbistatin | 1 uM, 10 uM |
| Cadmium chloride | 1 mM, 5 mM, 10 mM |
| Ciliobrevin-a | 100 uM |
| Ciliobrevin-d | 100 uM |
| Cobalt chloride | 1 mM, 5mM, 10 mM |
| Dibucaine | 1 mM, 10 mM |
| Gadolinium Chloride | 50 uM |
| Ionomycin | 1uM, 5 uM |
| Latrunculin a | 100 nM, 1 uM |
| Nickel(II) Chloride | 1 mM, 5 mM, 10 mM |
| Nocodazole | 0.1 ug/mL, 1 ug/mL |
| Potassium Chloride | 1 mM, 10 mM |
| Tetraethylammonium Bromide | 1 mM, 5 mM, 10 mM |
| Thapsigargin | 100 nM, 1 uM |
| Trifluoroperazine | 1 uM, 10 uM |
| Verapamil | 1 uM, 10 uM, 1 mM |
| Y-27632 | 10 uM, 50 uM |

